# Using CRISPR/Cas9 to identify genes required for mechanosensory neuron development and function

**DOI:** 10.1101/2023.05.08.539861

**Authors:** Christopher J. Johnson, Akhil Kulkarni, William J. Buxton, Tsz Y. Hui, Anusha Kayastha, Alwin A. Khoja, Joviane Leandre, Vanshika V. Mehta, Logan Ostrowski, Erica G. Pareizs, Rebecca L. Scotto, Vanesa Vargas, Raveena M. Vellingiri, Giulia Verzino, Rhea Vohra, Saurabh C. Wakade, Veronica M. Winkeljohn, Victoria M. Winkeljohn, Travis M. Rotterman, Alberto Stolfi

## Abstract

Tunicates are marine, non-vertebrate chordates that comprise the sister group to the vertebrates. Most tunicates have a biphasic lifecycle that alternates between a swimming larva and a sessile adult. Recent advances have shed light on the neural basis for the tunicate larva’s ability to sense a proper substrate for settlement and initiate metamorphosis. Work in the highly tractable laboratory model tunicate *Ciona robusta* suggests that sensory neurons embedded in the anterior papillae of transduce mechanosensory stimuli to trigger larval tail retraction and initiate the process of metamorphosis. Here, we take advantage of the low-cost and simplicity of *Ciona* by using tissue-specific CRISPR/Cas9-mediated mutagenesis to screen for genes potentially involved in mechanosensation and metamorphosis, in the context of an undergraduate “capstone” research course. This small screen revealed at least one gene, *Vamp1/2/3*, that appears crucial for the ability of the papillae to trigger metamorphosis. We also provide step-by-step protocols and tutorials associated with this course, in the hope that it might be replicated in similar CRISPR-based laboratory courses wherever *Ciona* are available.

## Introduction

Solitary tunicates (*Ciona spp*.) have emerged as highly tractable model organisms for developmental, cell, and molecular biology (Cota 2018; Lemaire 2011). Tissue-specific CRISPR/Cas9-mediated mutagenesis has been adapted to *Ciona robusta* and is now routinely employed to test the functions of genes in *Ciona* embryos and larvae (Gandhi et al. 2018; Sasakura and Horie 2023). The low-cost and ease of CRISPR/Cas9 in *Ciona* makes these animals an ideal organisms for laboratory courses in higher education. Hands-on experience in CRISPR/Cas9 might prepare students for a world in which CRISPR/Cas9-based technologies become more prevalent (Thurtle-Schmidt and Lo 2018).

Here we used *Ciona robusta* in the context of an undergraduate “capstone” research course on the use of CRISPR/Cas9 in neurobiology, taught at the Georgia Institute of Technology. In this course, students selected 4 target genes from a list of genes putatively expressed in the mechanosensory neurons of the anterior papillae of the *Ciona* larvae. The papillae are a group of three small clusters of cells organized in a triangle at the anterior end of the larval head (**Figure 1**). Basic characterization of the cell types contained in these papillae suggest multiple adhesive, contractile, and sensory functions supporting the attachment of the larvae to the substrate and triggering the onset of metamorphosis (Nakayama-Ishimura et al. 2009; Zeng et al. 2019a; Zeng et al. 2019b). Recently, mechanical stimulus of the papillae was shown to be sufficient for triggering tail retraction, the first stage of metamorphosis (Wakai et al. 2021). This ability was shown to depend on PKD2-expressing papilla neurons specified by the transcription factor Pou4 (Sakamoto et al. 2022).

**Figure 1.**
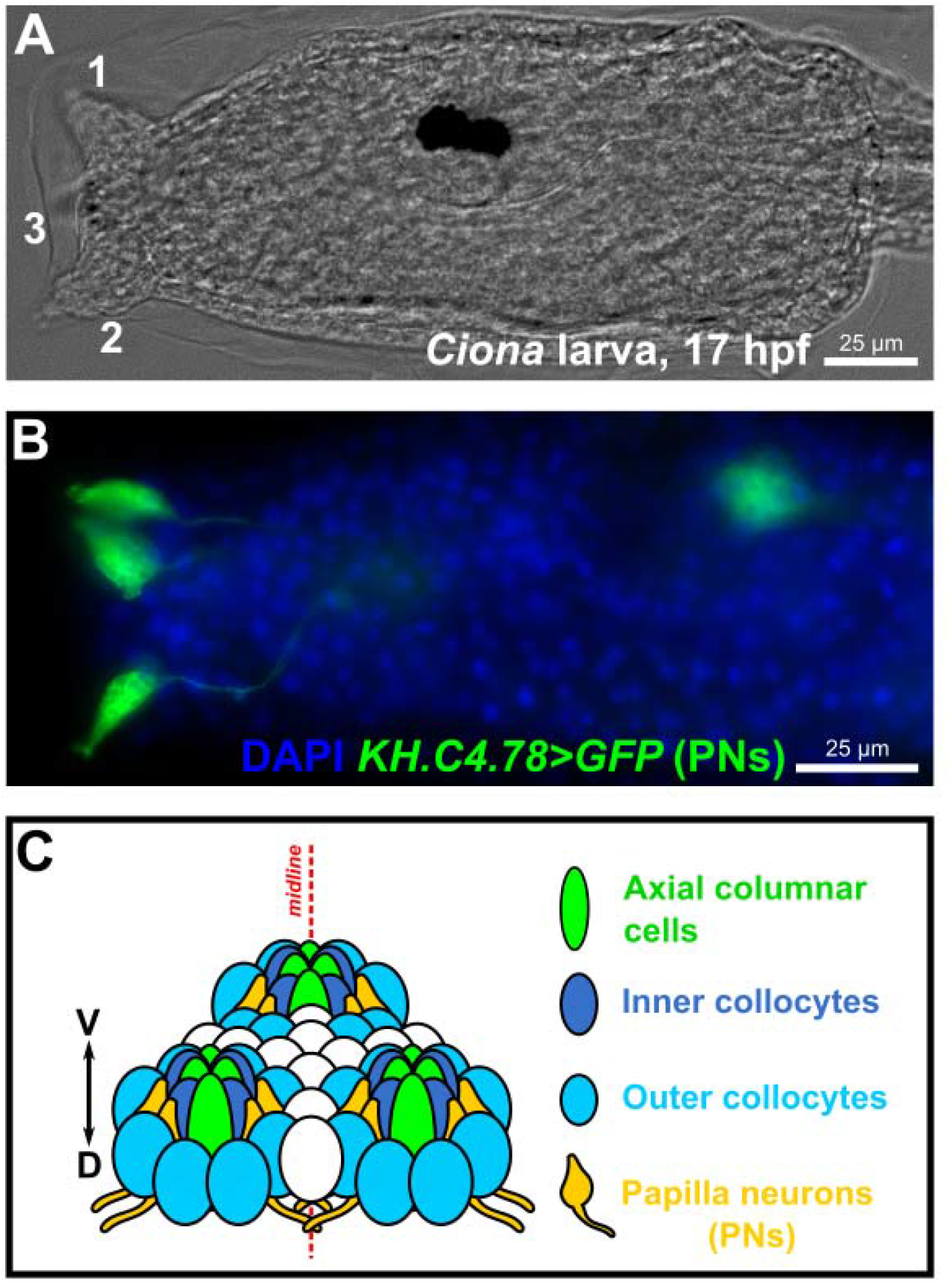
The sensory/adhesive papillae of the *Ciona* larva. Brightfield image of a *Ciona robusta (intestinalis* Type “A”) larva at 17 hours post-fertilization (hpf) raised at 20°C, showing the three protruding papillae of the head (numbered 1-3). Papilla number 3, the medial/ventral papilla, is out of focus. B) Image of electroporated *Ciona* larva at 17 hpf/20°C, papilla neurons (PNs) labeled by the reporter plasmid *KH.C4.78>Unc-76::GFP* (green, from Johnson et al. 2023). Nuclei counterstained by DAPI (blue). C) Summary diagram of the arrangement and cell type diversity of the papillae (from Johnson et al. 2023).

With this in mind, students in the course hypothesized that one or more genes expressed in the papillae neurons might be required for tail retraction and metamorphosis. Students designed and validated single-chain guide RNAs (sgRNAs) targeting four selected genes: *Tyrosine hydroxylase (TH), Vamp1/2/3, Neuronal calcium sensor 1 (NCS1)*, and *NARS1*. Of these, *Vamp1/2/3* was the only gene that, when knocked out, resulted in a metamorphosis defect.However, *NARS1* knockout in the developing central nervous system resulted in major morphological defects, indicating that our validated sgRNAs might still be instrumental in revealing the roles of these genes in other contexts. Here we describe our findings, in addition to providing detailed sequence information and protocols. We hope that this study will help other instructors who wish to implement a similar lab course based on CRISPR and/or *Ciona*, or researchers who wish to knock out these same *Ciona* genes out in other cell types.

## Methods

### Ciona handling, fixing, staining, and imaging

*Ciona robusta (intestinalis Type A)* were collected by and shipped from San Diego, CA (M-REP). Eggs were fertilized, dechorionated, and electroporated according to published protocols (Christiaen et al. 2009a; 2009b). Embryos were raised at 20°C. Embryos, larvae, and/or juveniles were fixed in MEM-FA solution (3.7% formaldehyde, 0.1 M MOPS pH 7.4, 0.5 M NaCl,1 mM EGTA, 2 mM MgSO4, 0.1% Triton-X100), rinsed in 1X PBS, 0.4% Triton-X100, 50 mM NH4Cl for autofluorescence quenching, and a final 1X PBS, 0.1% Triton-X100 wash. Specimens were imaged on a Leica DMI8 or Nikon Ti2-U inverted epifluorescence microscope.

### Phylogenetic trees

Protein sequences were aligned using online MAFFT version 7 (Katoh et al. 2019).Phylogenetic trees were assembled in MAFFT also, using default parameters: NJ (conserved sites), JTT substitution model, with heterogeneity among sites ignored (α = infinite) and no bootstrapping. Trees were visualized in MAFFT using Archaeopteryx.js (https://github.com/cmzmasek/archaeopteryx-js). Protein domain analysis was performed using SMART (http://smart.embl-heidelberg.de/)(Letunic et al. 2021).

### CRISPR/Cas9 sgRNA design and validation

Single-chain guide RNA (sgRNA) templates were designed using CRISPOR (Haeussler et al. 2016)(crispor.tefor.net) and synthesized custom-cloned into the U6>sgRNA-F+E vector (Stolfi et al. 2014) by Twist Bioscience (South San Francisco, CA). High Doench ‘16 score, high MIT specificity scores were prioritized, and targets containing known single-nucleotide polymorphisms were avoided. Validation of sgRNAs was performed by co-electroporating 25 μg of *Eef1a>Cas9* (Stolfi et al. 2014) and and 75 μg of the sgRNA plasmid, per 700 μl of total electroporation volume. Genomic DNA was extracted from larvae electroporated with a given sgRNA using a QIAamp DNA micro kit (Qiagen). PCR products spanning each target site were amplified from the genomic DNA, with each amplicon 150-450 bp in size. Amplicons were purified using a QIAquick PCR purification kit (Qiagen) and Illumina-sequenced using Amplicon-EZ service from Azenta/Genewiz (New Jersey, USA). Papilla-specific CRISPR knockouts were performed using *Foxc>Cas9*, as previously described (Johnson et al. 2023)

*Sox1/2/3>Cas9::GemininN* was constructed using the *Sox1/2/3* promoter (Stolfi et al. 2014) and the *Cas9::GemininN* as previously published (Johnson et al. 2023; Song et al. 2022). All sgRNA and primer sequences can be found in the **Supplemental Sequences File**. Detailed tutorials and protocols used for classroom activities can be found at the OSF link: https://osf.io/3fh89/Please contact the corresponding author to inquire about more detailed modifications to commercial kit manufacturers’ protocols.

## Results

### Selecting genes and designing sgRNAs

Genes to be targeted by CRISPR/Cas9 were chosen based on student preference, from a list of transcripts enriched in a cell cluster potentially representing the papilla mechanosensory neurons, identified from whole-larva singe-cell RNA sequencing data. Briefly, published data (Cao et al. 2019) were reanalyzed (Johnson et al. 2023) and papilla neuron identity was tentatively confirmed by enrichment with *Thymosin beta-related (KH*.*C2*.*140), Celf3/4/5 (KH*.*C6*.*128), Foxg (KH*.*C8*.*774), Synaptotagmin (KH*.*C2*.*101), Pou4 (KH*.*C2*.*42), Pkd2 (KH*.*C9*.*319), and TGFB (KH*.*C3*.*724)* based on previous reports (Horie et al. 2018; Katsuyama et al. 2002; Razy-Krajka et al. 2014; Sakamoto et al. 2022; Sharma et al. 2019; Zeng et al. 2019b)(**Supplemental Table 1**).To be clear, these are distinct from what we previously called “palp neurons” (Sharma et al. 2019), which were later identified conclusively as a non-neuronal cell type, the Axial Columnar Cells of the papillae (Johnson et al. 2020; Zeng et al. 2019b). The genes and sgRNAs selected for this study are detailed below.

### Tyrosine hydroxylase (KH.C2.252)

The gene selected by the first group of students was *Tyrosine hydroxylase (TH;* KyotoHoya gene model ID: *KH.C2.252)*, encoding the *C. robusta* ortholog of the rate-limiting enzyme of dopamine biosynthesis (Moret et al. 2005). Previously, *TH* was reported to be a marker of putative dopamine-releasing coronet cells of the ventral larval brain vesicle (Moret et al. 2005; Razy-Krajka et al. 2012; Takamura et al. 2010). Dopamine immunoreactivity was also observed in the papilla region of another species, *Phallusia mammilata* (Zega et al. 2005).

Pharmacological treatments suggested roles for dopamine in neuromodulation of larval swimming behavior in *Ciona* (Razy-Krajka et al. 2012), and suppression of metamorphosis in *P. mammillata* (Zega et al. 2005)Three sgRNAs were selected from those predicted by the web-based CRISPOR prediction tool (crispor.tefor.net)(Haeussler et al. 2016), as described in detail in the methods section and online protocols. Two were predicted to cut in exon 4 (named “TH.4.114” and “TH.4.140”) and one in exon 5 (“TH.5.44”)(**Figure 2A**). Because exon 5 encodes the beginning of the major catalytic domain of TH, these sgRNAs were predicted to generate frameshift mutations resulting in truncated proteins lacking the catalytic domain.

**Figure 2.**
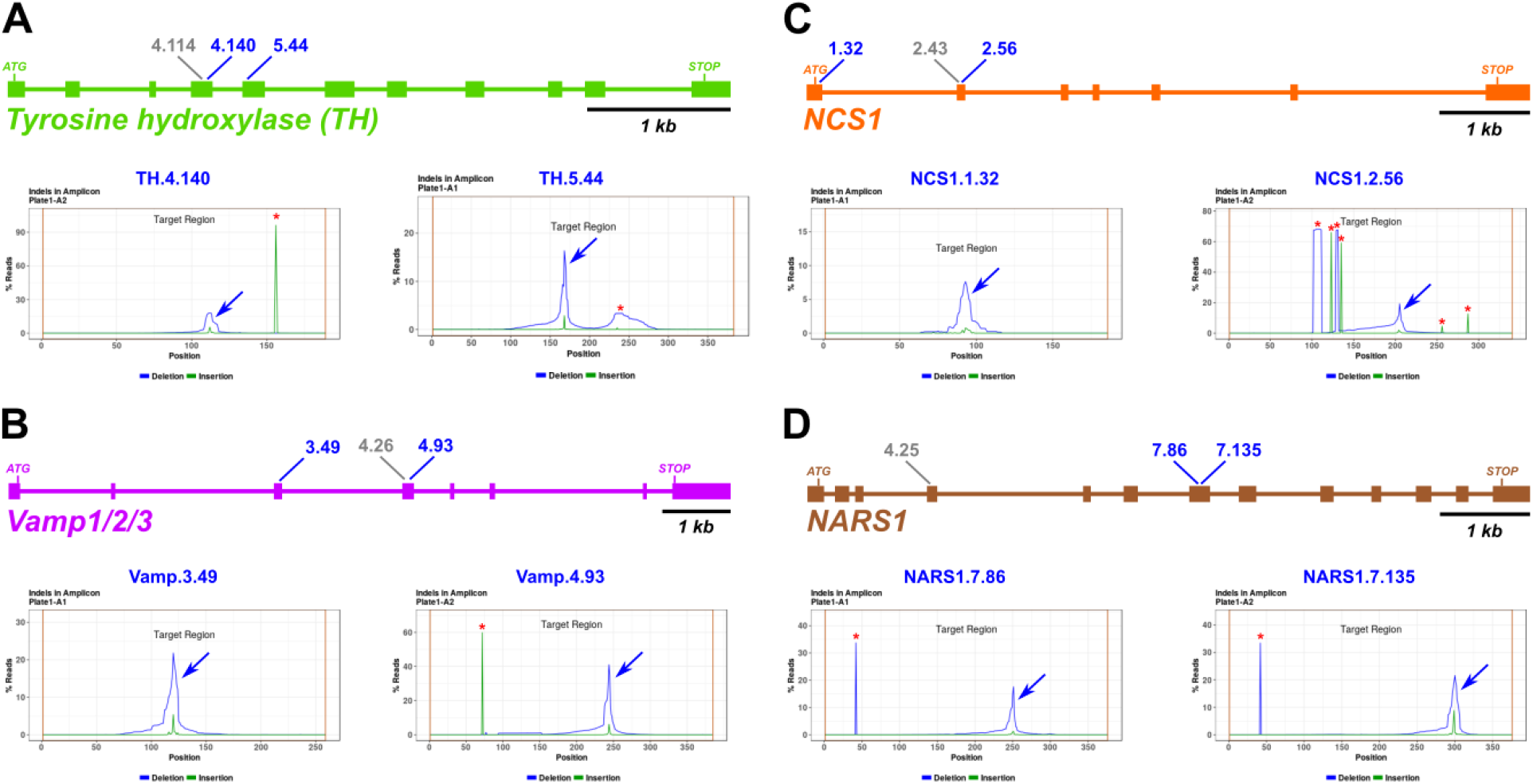
Design and validation of sgRNAs for CRISPR/Cas9-mediated mutagenesis. A-D) Diagrams of selected candidate gene loci and indel analysis plot for each selected sgRNA, based on next-generation sequencing of amplicons. Blue arrows indicate CRISPR/Cas9-induced indel peak, red asterisks indicate naturally-occurring indels. Blue sgRNA identifiers indicate top sgRNAs selected for phenotypic assay. Grey identifiers indicate sgRNA designed, tested, but not selected for further use.

### Vamp1/2/3 (KH.C1.165)

The second student group picked *Vamp1/2/3 (KH.C1.165)*, which encodes a member of the synaptobrevin family of SNARE complex proteins that carry out neurotransmitter vesicle release (Rizo 2022). Based on phylogenetic analysis in MAFFT (see methods), *Vamp1/2/3 (KH.C1.165)* appears to be orthologous to *VAMP1, VAMP2*, and *VAMP3* in humans (**Figure S1**). Its potentially evolutionarily conserved function and broad expression in the *Ciona* larval nervous system suggested an important role for *Vamp1/2/3* in neurotransmitter release in *Ciona*, including in the papilla neurons during settlement. The *Vamp1/2/3* gene in *Ciona* appears to give rise to a few different alternatively spliced isoforms. The sgRNAs selected from CRISPOR included one sgRNA targeting exon 3 (“Vamp.3.49”) and two sgRNAs targeting exon 4 (“Vamp.4.26” and “Vamp.4.93”) in the “v3” and “v4” transcript variants (**Figure 2B**). These exons become exons 2 and 3, respectively, in all other transcript variants.

### Neuronal calcium sensor 1 (KH.C1.1067)

Group number 3 selected the gene *Neuronal calcium sensor 1 (NCS1*, gene model *KH.C1.1067)*. According to our phylogenetic analysis, KH.C1.1067 appeared to be most similar to human NCS1 and its *Drosophila melanogaster* orthologs, Frequenin1 and Frequenin2 within the NCS family of proteins (**Figure S2A**). NCS1/Frq proteins regulate neurotransmission through both pre- and post-synaptic mechanisms (Dason et al. 2012), likely on account of their ability to bind Ca2+ ions through their multiple EF hand domains. In *Ciona, NCS1* had been previously identified as a transcriptional target of Neurogenin in the Bipolar Tail Neurons of the larva, suggesting a broader role in neuronal function (Kim et al. 2020). However, no function has yet been shown for this gene in *Ciona*. Three sgRNAs targeting *NCS1* were selected for testing: one sgRNA targeting exon 1 (“NCS1.1.32”) and two sgRNAs targeting exon 2 (“NCS1.2.43” and “NCS1.2.56”)(**Figure 2C**). As these sgRNAs are predicted to cut 5’to the exons encoding the EF hand domains (exons 3-7, **Figure S2B**), the resulting frameshift mutations are predicted to result in a truncated, non-functional polypeptide.

### NARS1 (KH.C12.45)

The fourth student group picked *NARS1 (KH.C12.45)*, which encodes the *C. robusta* ortholog of Aparaginyl tRNA synthetase 1 (cytoplasmic), which catalyzes the attachment of asparagine (Asn/N) to its cognate tRNAs (Shiba et al. 1998). In a neurodevelopmental context, it has been shown that loss of *NARS1* in human brain organoids impairs neural progenitor proliferation (Wang et al. 2020). Mutations in *NARS1* is associated with various neurodevelopmental syndromes such as microcephaly and cognitive delays (Wang et al. 2020), suggesting that regulation of protein synthesis rates is indispensable for development of the nervous system. Phylogenetic analysis shows that these aminoacyl-tRNA synthetases are highly conserved in their specificity, with simple 1-to-1 orthology between *Ciona* and human genes of various types and classes within this gene family (**Figure S3A**). For this gene, one sgRNA targeting exon 4 (“NARS1.4.25”) and two sgRNAs targeting exon 7 (“NARS1.7.86” and “NARS1.7.135”) were designed (**Figure 2D**). While the tRNA anti-codon domain is encoded by exons 5-7, and the tRNA synthetase domain is encoded by exons 7-13, these sgRNAs are predicted to result in truncated NARS1 polypeptides lacking both major functional domains (**Figure S3B**).

### Validation of sgRNA efficacy by Illumina amplicon sequencing

Validation of sgRNA efficacies was performed by sequencing amplicons surrounding each target site, from larvae electroporated with a given sgRNA vector together with the ubiquitously-expressed *Eef1a>Cas9* (Stolfi et al. 2014)(**Figure 3A**). Although we had previously reported a Sanger sequencing-based method for estimating mutagenesis efficacy (Gandhi et al. 2018), that strategy is frequently hampered by naturally-occurring indels and poor sequencing quality. We decided instead to quantify mutagenesis by sequencing amplicons using a commercially available Illumina-sequencing based service, as recently described (Johnson et al. 2023).

**Figure 3.**
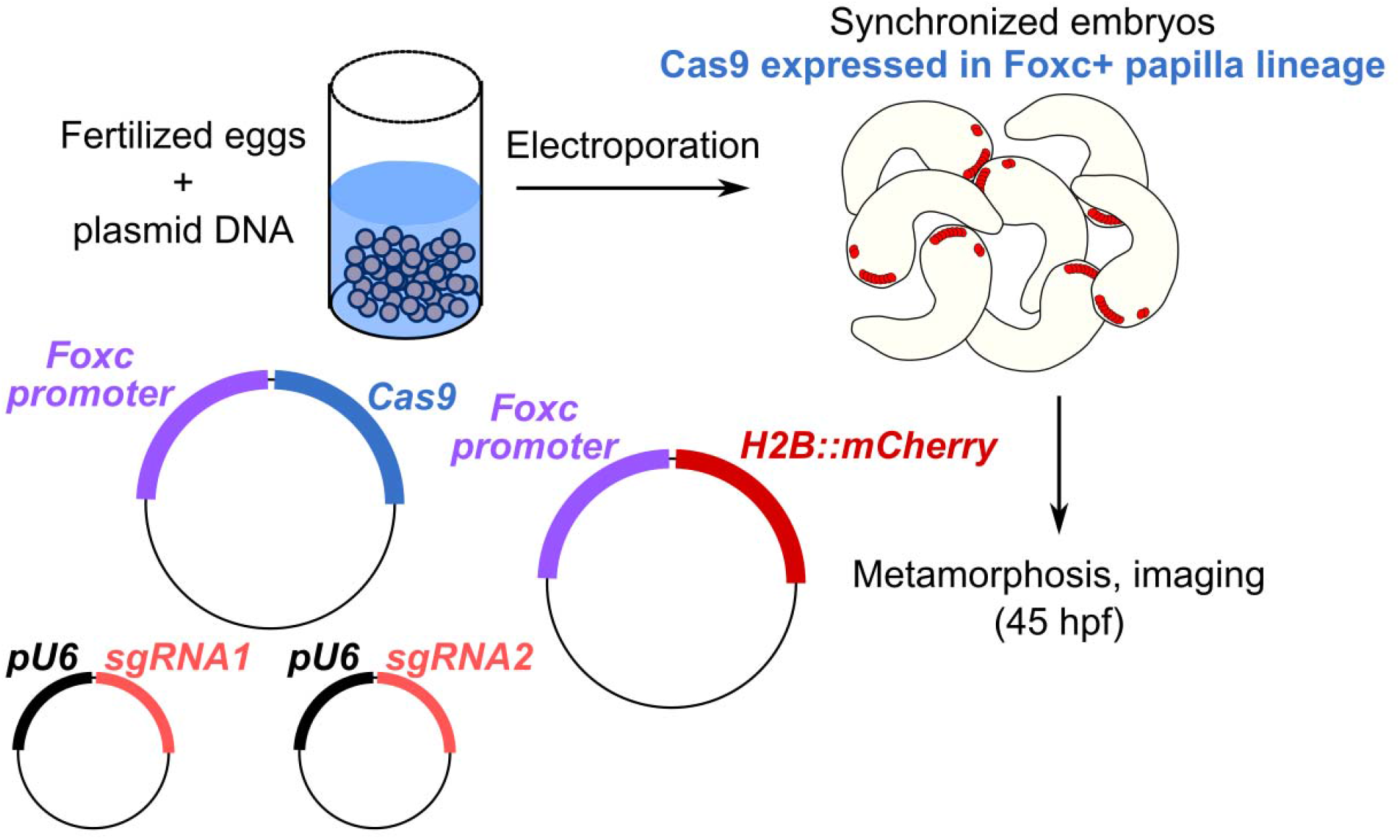
Tissue-specific CRISPR/Cas9-mediated mutagenesis for tail retraction assay.

Briefly, 75 μg/700 μl total electroporation volume of each sgRNA plasmid was co-electroporated with 25 μg/700 μl of *Eef1a>Cas9* into zygotes, which were collected at larval stage (∼17 hours post-fertilization, reared at 20°C). Larvae electroporated with the same sgRNA vector were pooled, and genomic DNA extracted from them. DNA fragments spanning each target site, ranging from 150-450 bp as required by the sequencing service, were amplified by PCR from each genomic DNA pool. Negative controls for each amplicon were derived by repeating the PCR on samples from larvae electroporated with sgRNA vector targeting a different sequence (e.g. targeting exon 2 instead of exon 3). Amplicons were then submitted for library preparation, sequencing, and analysis by Genewiz/Azenta. The sgRNAs chosen for further experiments were those that resulted in a larger portion of on-target indels based on visual examination of the indel plot automatically generated by the amplicon sequencing service. This was better than relying on raw mutagenesis rates provided by the service, as the plots revealed a high frequency of naturally occurring indels that prevented the automatic quantification of the true efficacies of some sgRNAs. This new approach is described in greater detail in the methods and online protocols.

Amplicon sequencing revealed the indels induced by all sgRNAs except for NARS1.4.25, which was not evaluated due to failure to amplify its target site by PCR (**Figure 2, S4**). For *TH*, we found that all three sgRNAs were effective at generating indels at the target sequences, though TH.4.140 and TH.5.44 were selected, as having barely edged out TH.4.114 (**Figure 2A, S4A**). For *Vamp1/2/3*, the most efficient sgRNAs was Vamp.4.93 (>40% efficacy, **Figure 2B**), while Vamp.3.49 and Vamp.4.26 were less efficacious at ∼20-30% indels (**Figure S4B**). Because we wished to use a pair of sgRNAs targeting different exons, we selected Vamp.4.93 and Vamp.3.49 for further use. All three sgRNAs targeting *NCS1* generated indels, though NCS1.1.32 efficacy was only 12% indels (**Figure 2C, S4C**). Because NCS1.2.43 and NCS1.2.56 targets overlapped, we paired the most efficacious sgRNA (NCS1.2.56) with NCS1.1.32. Finally, NARS1.7.86 and NARS1.7.135 resulted in mutagenesis efficacy rates >15% (**Figure 2D**). As these were the only two *NARS1-*targeting sgRNAs for which amplicons were successfully amplified by PCR (**Figure S4D**), we proceeded with both and did not further use the untested sgRNA NARS1.4.25.

### Papilla lineage-specific knockout of target genes by CRISPR/Cas9

It was recently shown that knockdown or knockout of the neuronal transcription factor *Pou4* eliminates papilla neuron formation and subsequently, papilla neuron-induced signals for tail retraction and metamorphosis (Sakamoto et al. 2022). Other CRISPR gene knockouts were shown to result in a mixture of tail retraction and body rotation defects during settlement and metamorphosis (Johnson et al. 2023). We therefore used papilla-specific CRISPR/Cas9-mediated knockout in F0 embryos to test the requirement of our candidate genes in a similar tail retraction assay. We used the *Foxc* promoter (Wagner and Levine 2012) to drive expression of Cas9 in the anteriormost cells of the neural plate, which gives rise to the entire papilla territory and part of the oral siphon primordium (**Figure 3**). Embryos were electroporated with 40 μg/700 μl *Foxc>Cas9*, 10 μg/700 μl *Foxc>H2B::mCherry*, and gene-specific pairs of sgRNA vectors (40 μg/700 μl each sgRNA vector). “Positive control” embryos were electroporated as above, using a previously published pair of sgRNA vectors targeting *Pou4* (Johnson et al. 2023), and “negative control” embryos were electroporated with 10 μg/700 μl *Foxc>H2B::mCherry* alone.

All embryos were raised through larval hatching and settlement, and fixed at 45 hours post-fertilization, upon which tail retraction and body rotation were scored (**Figure 4A**). As previously reported, *Pou4* knockout in the papilla territory resulted in frequent block of tail resorption and body rotation compared to the negative control (**Figure 4B**). Of the gene-specific CRISPR samples, only *Vamp1/2/3* CRISPR showed a substantial effect on metamorphosis, with only 56% of H2B::mCherry+ individuals having retracted their tails. This was closer to the *Pou4* CRISPR (20% tail retraction) than to the negative control (97% tail retraction). The effect of *Vamp1/2/3* CRISPR on body rotation was very similar (**Figure 4B**). An independent replicate of *Vamp1/2/3* CRISPR confirmed this result (**Figure S5**). Taken together, these data suggest that knocking out *Vamp1/2/3* in the papilla territory impairs the ability of the larva to trigger the onset of metamorphosis.

**Figure 4.**
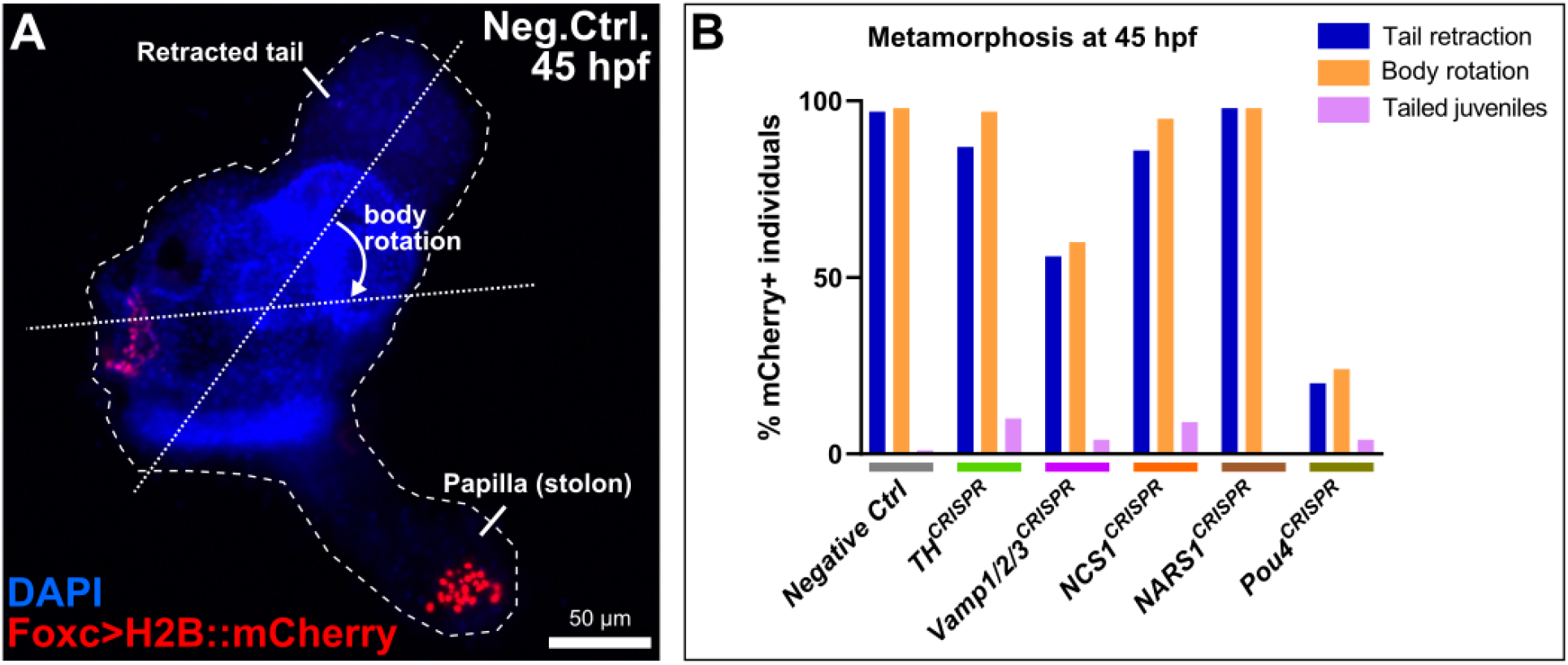
Scoring metamorphosis defects in CRISPR larvae. A) Example of a juvenile at 45 hours post-fertilization (hpf) in the “negative control” population, showing the retracted tail and body rotation that occurs during metamorphosis. *Foxc>H2B::mCherry* (red) labels the cells of the oral siphon and the papillae, the latter of which are transformed into the stolon of the juvenile. Nuclei counterstained by DAPI (blue). B) Scoring of *Foxc>H2B::mCherry+* individuals upon papilla-specific CRISPR/Cas9-mediated mutagenesis of the selected candidate genes. “Tailed juveniles” are individuals that have undergone body rotation but not tail retraction. *Pou4* CRISPR served as the “positive control”, eliminating the papilla neurons that trigger metamorphosis (see text for citations). Of the four genes tested, only *Vamp1/2/3* CRISPR appeared to result in substantial loss of tail retraction and body rotation, though not as penetrant as the *Pou4* CRISPR.

### Neural tube-specific knockout of *NARS1* causes neurulation defects

Because *NARS1* is associated with various neurodevelopmental defects in mammals (Wang et al. 2020), NARS1 students also tested the requirement of *NARS1* in *Ciona* neurulation. *NARS1* was targeted in the neurectoderm using *Sox1/2/3>Cas9::GemininN*, and the *Nut>Unc-76::GFP* reporter was used to visualize the central nervous system (Shimai et al. 2010). Embryos were electroporated with 40 μg/700 μl *Sox1/2/3>Cas9::GemininN*, 40 μg/700 μl *Nut>Unc-76::GFP*, and 40 μg/700 μl each of both *NARS1* sgRNA vectors. As a result of *NARS1* CRISPR in the neurectoderm, a high frequency of curled/twisting tails specifically in the CRISPR larvae, but not in the negative control (**Figure S6**). This was scored as well, and curved tails were observed in 24 out of 50 *NARS1* CRISPR larvae (48%), compared to 0 out of 50 negative control larvae (0%). Upwards curvature of the tail is a hallmark of impaired neural tube closure in *Ciona* (Mita and Fujiwara 2007), suggesting *NARS1* may be required in the neural tube for proper neurulation.

### Conclusion

We have described the design and validation of sgRNAs targeting four different genes in *Ciona*, in the context of a university-level laboratory course. Of these, only one gene (*Vamp1/2/3*) was shown to be required for tail retraction and body rotation at the onset of metamorphosis. Although the other CRISPR knockouts did not result in a noticeable metamorphosis defect, our validated sgRNAs may be of great interest to other *Ciona* researchers studying these genes in other contexts.

Our results do not entirely rule out a role for the other three genes tested. For instance, there may be similar genes with overlapping functions that can compensate for the loss of one of them. In fact, another NCS family gene, *KH.C9.113*, was also found to be enriched in the putative papilla neuron cell cluster by scRNAseq (**Supplemental Table 1**). Another possibility is that the gene may be required for fine-tuned mechanosensory discernment of settlement substrates in the wild, while in our laboratory assays most larvae eventually retract their tails as long as the papilla neurons retain most of their functions.

The requirement of *Vamp1/2/3* for papilla neuron-mediated tail retraction is not surprising, given its central role in synaptic transmission. However, our results and methods described here establish a proof-of-principle for future screens for genes potentially important for mechanosensory papilla neuron development and function in *Ciona* larvae.

## Supporting information

Supplemental Sequences File

## Acknowledgments

We thank Dexter Dean and Alison Onstine for managing the Neuroscience undergraduate teaching lab and granting access to equipment and materials for the course. We thank the students in the previous iteration of this course for their constructive feedback and suggested improvements to the teaching material. We thank Lindsey Cohen for technical assistance. This work was supported by Georgia Tech student fees and institutional funds, an NSF GRFP award to CJJ, NIH award K99NS126576 to TMR, NSF IOS award 1940743 to AS, and NIH award R01GM143326 to AS.

---

**Figure S1.**
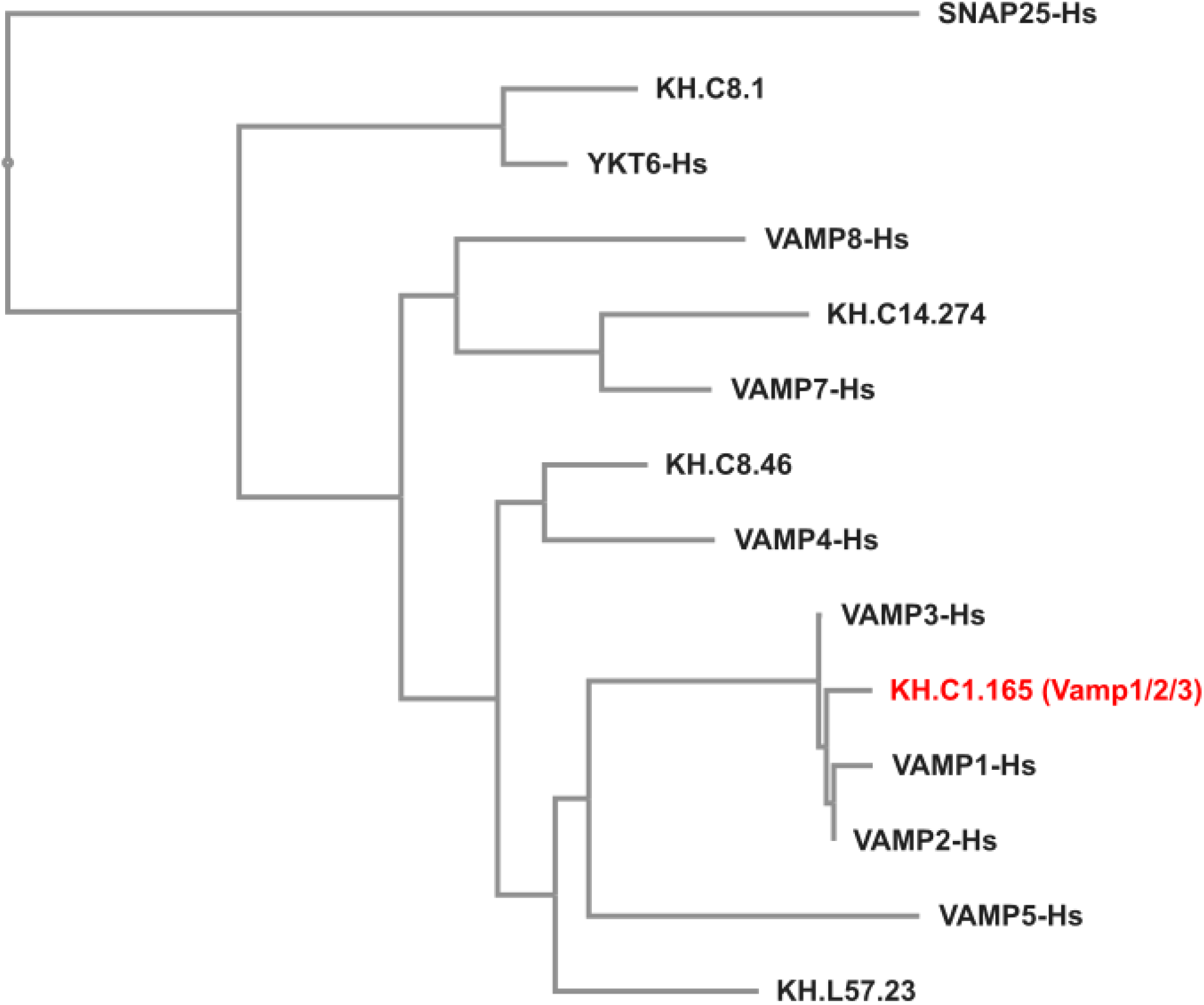
Phylogenetic tree of Vamp proteins. Tree showing phylogenetic analysis of predicted proteins encoded by *VAMP* family genes from human (Hs) and *Ciona robusta* (KH gene models). See methods for details and supplemental sequences file for protein sequences used.

**Figure S2.**
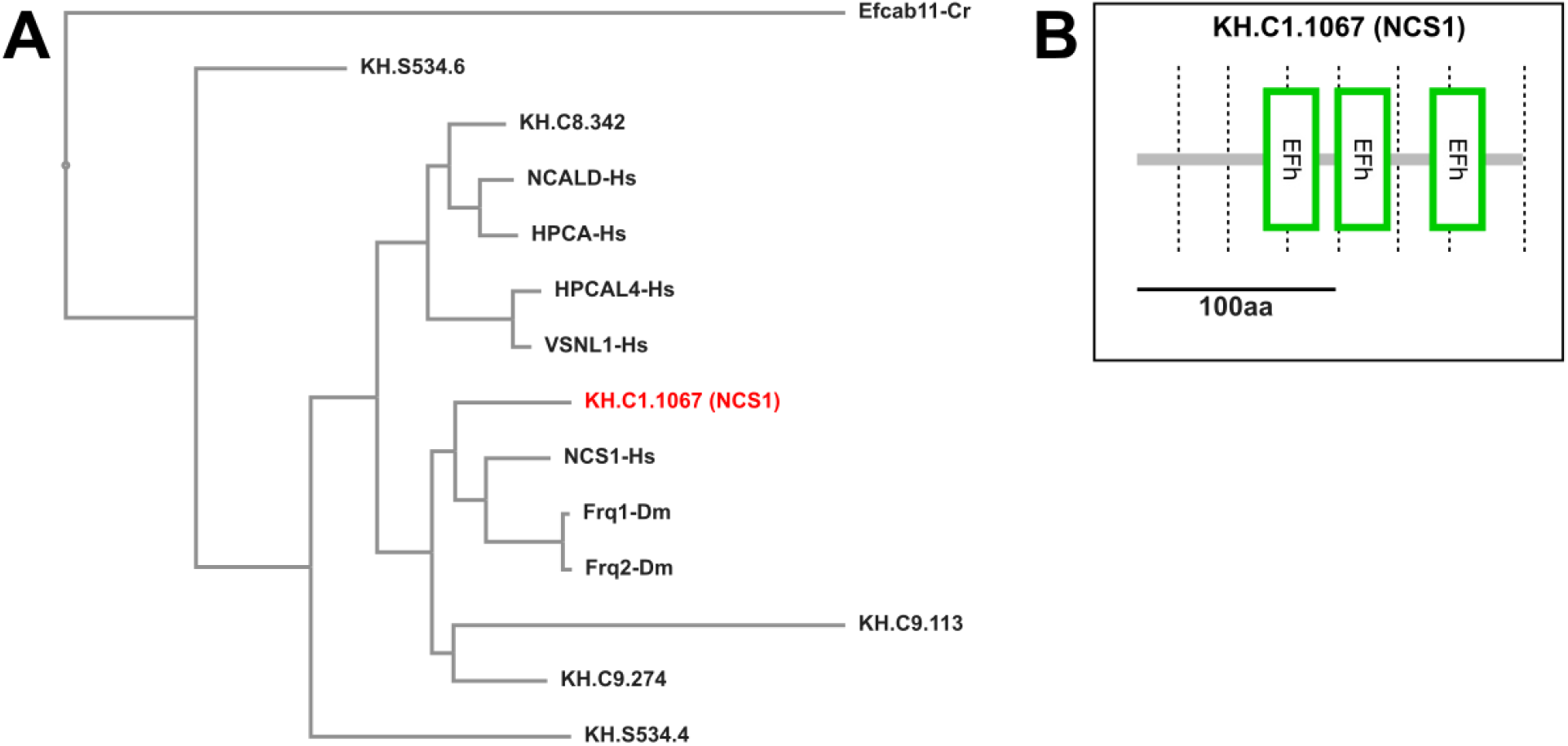
Phylogenetic tree of NCS proteins and diagram of *Ciona robusta* NCS1 domains. A) Tree showing phylogenetic analysis of proteins encoded by *Neuronal Calcium Sensor (NCS)* family genes from human (Hs), *Drosophila melanogaster* (Dm) and *Ciona robusta* (KH gene models). Efcab11 from *C. robusta* (Cr) was used to root the tree. See methods for details and supplemental sequences file for protein sequences used. B) Protein domain analysis diagram of *Ciona robusta* NCS1 from SMART (Letunic et al. 2021) showing its predicted three EF-hand (Efh) domains. Dashed lines indicate exon-exon junctions.

**Figure S3.**
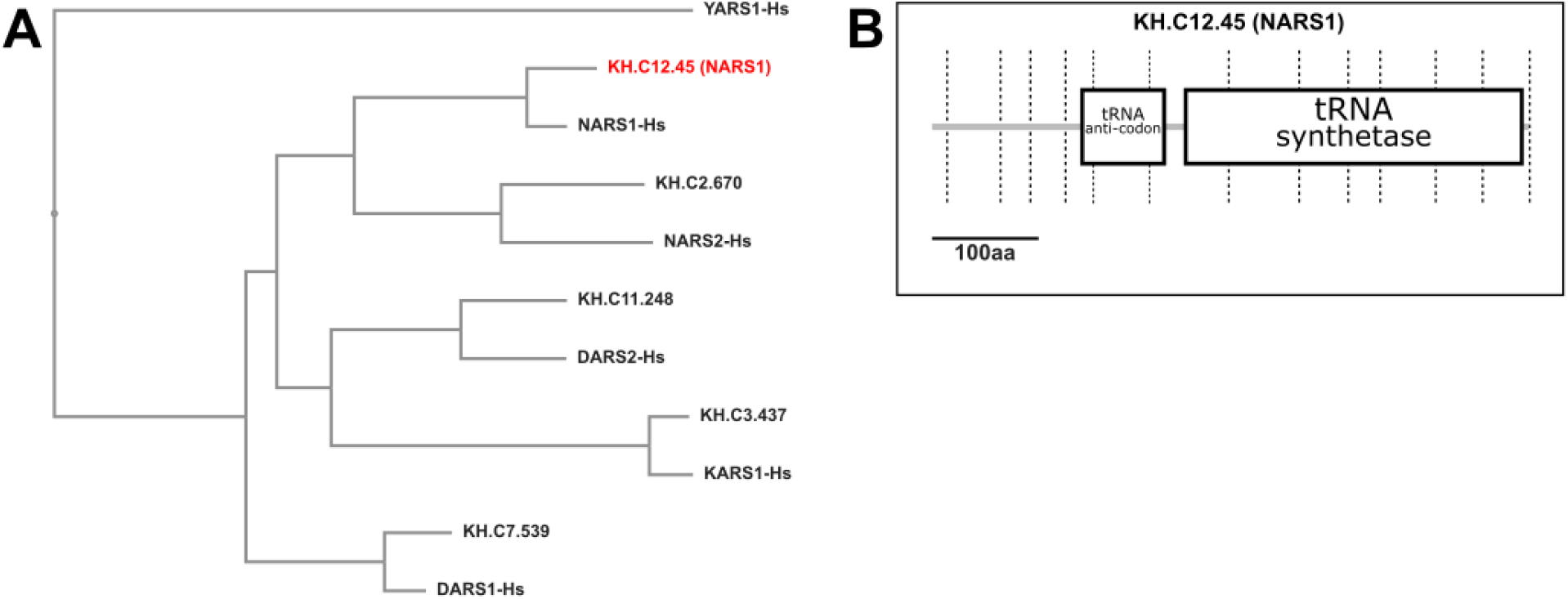
Phylogenetic tree of select aminoacyl-tRNA synthetases and *Ciona* NARS1 domains. A) Tree showing phylogenetic analysis of proteins encoded by a sampling of aminoacyl-tRNA synthetase genes from human (Hs) and *Ciona robusta* (KH gene models). See methods for details and supplemental sequences file for protein sequences used. B) Protein domain analysis diagram of *Ciona robusta* NARS1 from SMART (Letunic et al. 2021) showing its predicted domains. Dashed lines indicate exon-exon junctions.

**Figure S4.**
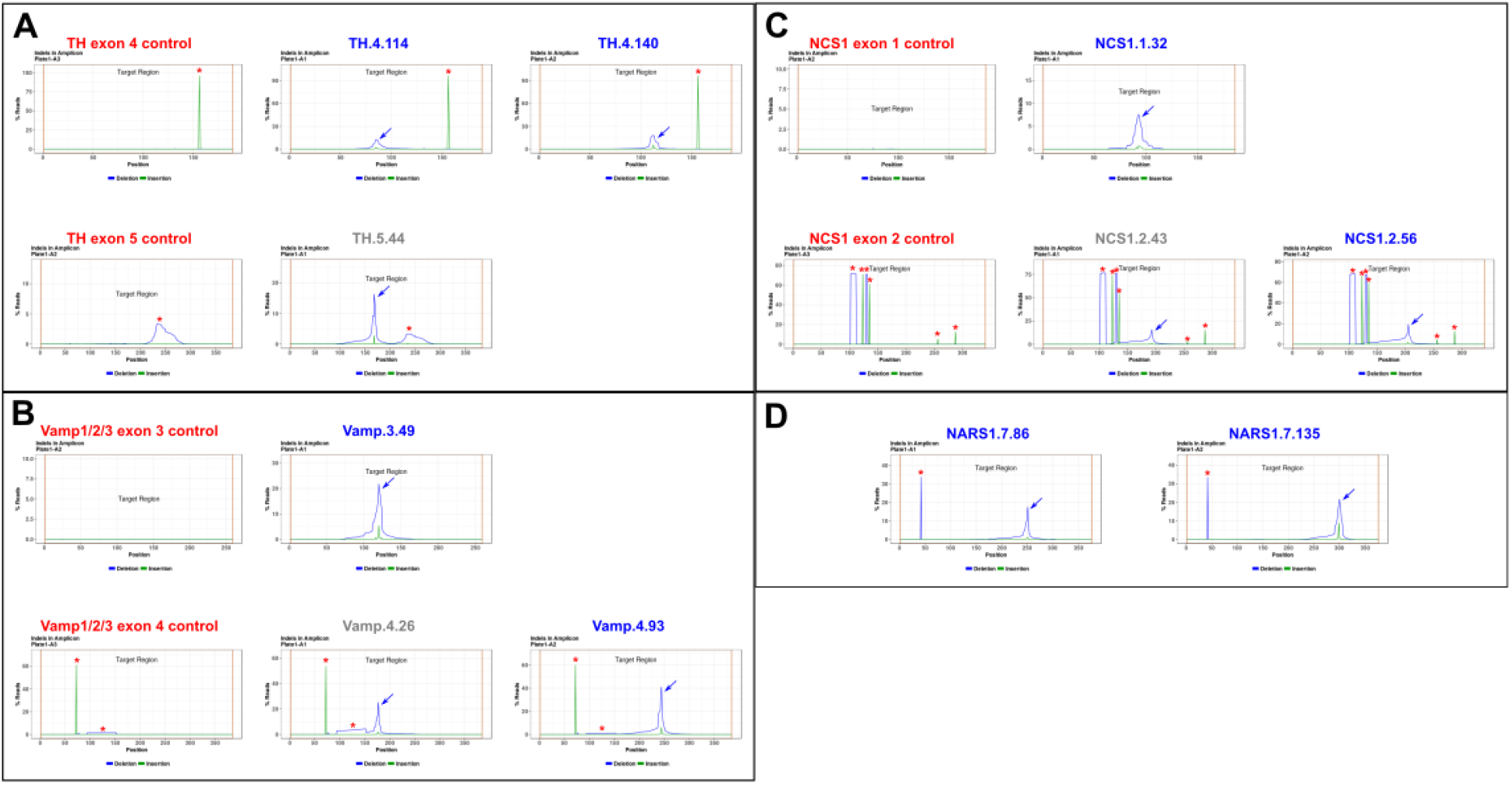
Indel plots for all sgRNAs tested. A-D) NGS indel validation plots (including negative controls) for all the sgRNAs tested in this study. No amplicon was obtained for the third *NARS1* sgRNA nor the *NARS1* negative control. Blue arrows indicate CRISPR-generated indels, red asterisks indicate naturally occurring indels.

**Figure S5.**
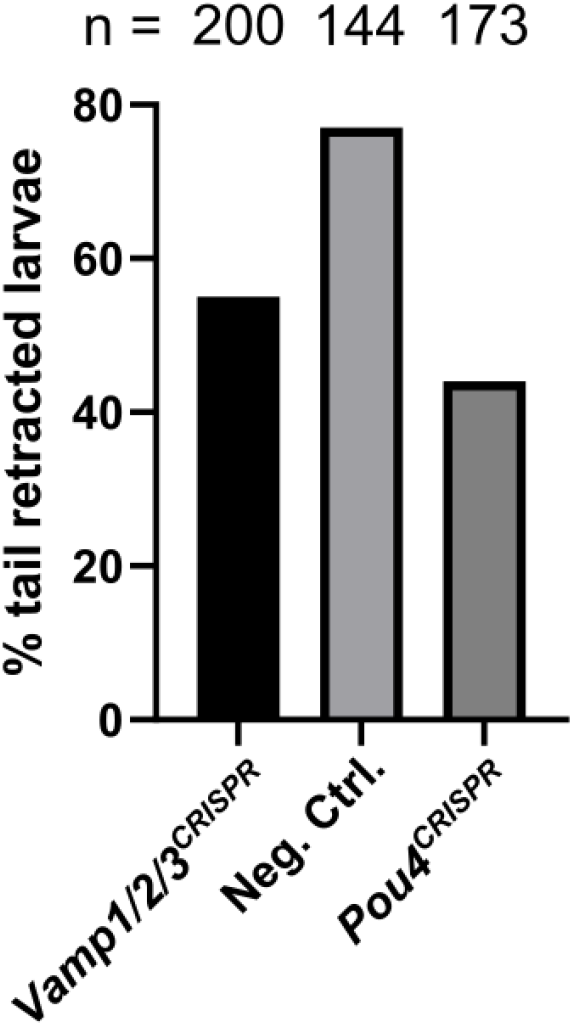
Replicate of *Vamp1/2/3* CRISPR. Independent replicate of papilla-specific *Vamp1/2/3* CRISPR in *Ciona* larvae. Embryos were electroporated with 40 μg/700 μl *Foxc>Cas9* and gene-specific pairs of sgRNA vectors (40 μg/700 μl each sgRNA vector). Negative control embryos were electroporated with 40 μg/700 μl *Foxc>Cas9* alone. Tail retraction was scored at 48 hours post-fertilization without screening for mCherry+ individuals.

**Figure S6.**
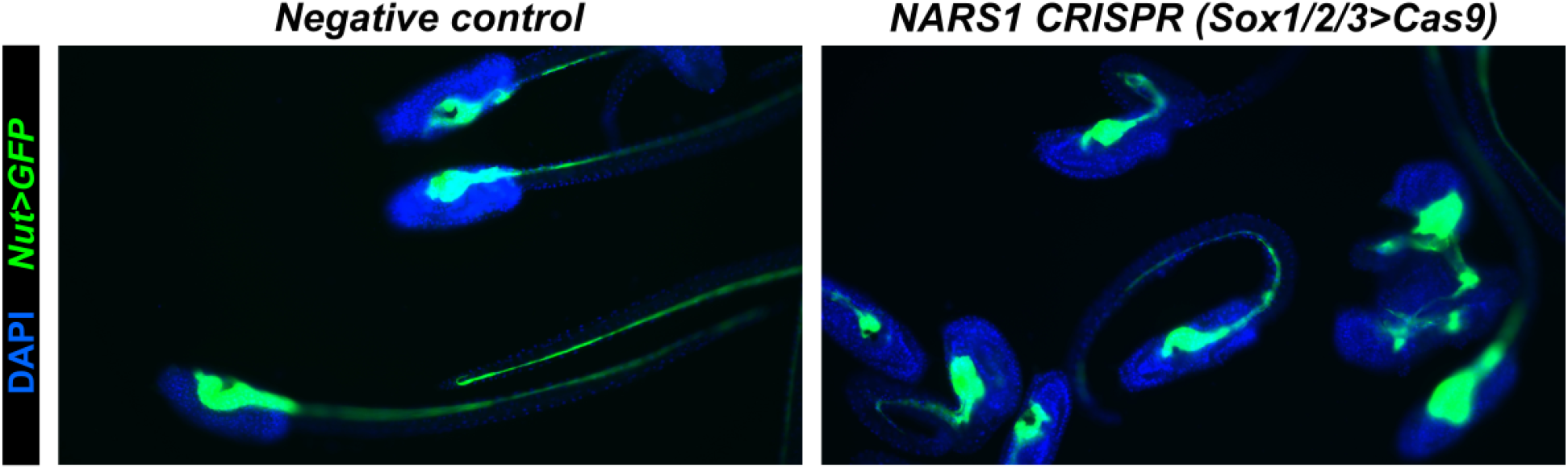
Neurectoderm-specific knockout of *NARS1* impairs neurulation. Negative control larvae showing normal neural tube and tail morphogenesis, compared to *NARS1* CRISPR larvae. *Nut>Unc-76::GFP* (green) labels the central nervous system. Nuclei counterstained by DAPI (blue). See text for quantification.

