## Supplemental Sequences File for "Using CRISPR/Cas9 to identify genes required for mechanosensory neuron development and function"

**Supplemental sequences file – Johnson et al. 2023 (Using CRISPR/Cas9 to...)**

>Nut -1155/-1 (Nut promoter based on Shimai et al. 2010)

(Sequence of the Nut driver based on genome assembly, not verified)

atctgttctaggaatcttttaactcggcgagcctgtgttttctttttagactataataccctccaataacagggtaagatgcatgcgaaaaaaacgattgattttgtgcatataaaatatttataatagtctggggttatgtgagtgaaatagctggttatatctaactgcagccggaaatacgtgctctttgcagaaaggaaaatccatttctcgtgtattgcaacatacggtatagtaggtttagatgggtcacctttaccacaaataacatccaaatatcctgagcatgttttaaacatttaacaacggtatatgtggaagttgcgagaatacggtttgataattcattaaatatgctttgtttactacaaaatgggacgggaaaataaaataaaaaggtgtcctatcttcccccatcctactatatatatcatattttctaatcctgttttagcaattaacaacgctcgtttaaagtcgtgggtatatgcggttacataaatttaaaaatattatttgtttacttcaaaatagaacgataaaagcgtgtataaacatgttgtatcatcttaccccatcctgctacaaatataaacaatttcatgataagaaaatgcacatgtttgttatgccgcgtgttgtgtacagttctggttaggatttcagaacacacacacagaaaatcggtcgtgcctgctgctgcaactagtgcgacacctcaataattatgagactgcctcagtgcattcctatcaacgtgttttgcgctcgtatcacccaaaaagcgtgcctcgtgctgcctacacggtgtggctacataaaaacgtcgcgtgttttactgttccagttcgtttttgtattcgtggaaataggttcgcattttttattaagaaagagtttggttttcgaggtcactgaattgcaattagaaggcatttaatagtagcactaggacactctattcactgcgttgtctaaatccatgcgaaacaacaaaataagtcgcaagacatgctgtgcgtgtattgtttttttgaaaccccgtcgctttgtgaaaatctggttgattattttttcgtaccagttttacagtttaaatacgactgtgcttcagtttttgttagtattgagttgtacactatatcaacaac

**sgRNAs used in this study:**

TH.4.114

**GCTGGAATCAATTGGTTGCA (G+N19)**

TH.4.140

**GAATTGTTAACGGGACTGAA (G+N19)**

TH.5.44

**GCTGAGCTTGAATTGTGCAG (G+N19)**

Vamp.3.49

**GGTCCAGCTGCACCAACTGG (G+N19)**

Vamp.4.26

**GGACATCATGAGGACAAATG (G+N19)**

Vamp.4.93

**GGGGCAGATGCATTACAGCA (G+N19)**

NCS1.1.32

**GGGCAGCAAACTGAGCGCAG (G+N19)**

NCS1.2.43

**GTGGTATCGAGACTTTATGA (G+N19)**

NCS1.2.56

**GTTATGAAGGATTGTCCCAA (G+N19)**

NARS1.4.25

**GAGAACAAGAAGATGCGGAG (G+N19)**

NARS1.7.86

**GATAGATACAGTCCTCAACC (G+N19)**

NARS1.7.135

**GACAGACATCTAGTTATCCG (G+N19)**

Pou4.3.21 (Johnson et al. 2023)

**GCTGAGTGGTGGAAAGCGGG (G+N19)**

Pou4.4.106 (Johnson et al. 2023)

**GAGGATGGAAATGATTCGGG (G+N19)**

*Primers for amplicon sequencing to measure mutagenesis efficacy*

| **Gene + exon** | **Forward primer** | **Reverse primer** |
| --- | --- | --- |
| TH exon 4 | ACACTAATTAAAAACGTTGCCG | CTATATGTGCCAATAGTGCCC |
| TH exon 5 | TGGGCACTATTGGCACATA | ACATAAATGACGGTAGAATACCC |
| Vamp1/2/3 exon 3 | GCACTTAAACCCTTATATCCCTG | CGATTTTTGACCCGAAGAACA |
| Vamp1/3/4 exon 4 | AGAAACCATGGTGTGCCTC | ATAAGCGTCCACCATTTCCC |
| NCS1 exon 1 | GTGATCTTTGTAGGTCTTCTTT | AGTTAGAAACTAAACGACCGC |
| NCS1 exon 2 | CTGGTTGCTCGCTATAAGTAA | CGTATTCTTGCGACTCTTATAGG |
| NARS1 exon 4 | GGCTACCCTGAATAATCAAAACTT | TTAGCCGTTGTACTGAAGTCC |
| NARS1 exon 7 | GTTGGTATGCCCCTAATTACC | CAAAATCACCTCATAATATCCTCG |

**Sequences for Vamp1/2/3 phylogenetic tree:**

>KH.C1.165.v3.A.SL2-1 (Vamp1/2/3)

MSGYDAVPNPRSYGGTSNYGGPAAPTGGSSQASKKLQQTQAQVDDVVDIMRTNVDKVLERDQKLSELDDRADALQQGASQFETQAAKLKRKYWWKNCKMWAILIIVILVIIIIIVVSVVTQTQKKT

>KH.C8.1.v3.A.SL2-2

MVKLFSLSLFYKGESCVIPLIAEYDTDSFSFFQRKSVQEFLSFTSKIITERTHPGMRQSVKEQEHLCHVYVREDRLAAVLISDEDYPRRVAFTVLTKVCEDFAKKFPPSELNNPAPMNIQFEGVREMLCKYQNPKEADSMSRVQAELDETKIVLHGTMESLLQRGEKLDDLVAKSDQLSSQSKMFYKQIKIHMKHDKHLWKKCTIAYRANFHLV

>KH.C14.274.v1.B.ND1-1

MTILFSIIARGTTVLARYASCAGNFQEVSEQILSKITADDAKLTYSHGSYLFHYICDDRIVYMAITEDDFERSKAFRYLSDIRKKFQSTYGKNVHTALPYAMDTDFARVLMVQMKRFSSNEEPENKVEEVQDQLNDLKGIMVKNIDSIANRGENLNLLVDKTEDLSESAVTFKKQSTTLRRRLWWKNVKITVILVIVAIIVLYFIICAACGGMNWPRCVHQSNSPNKTLTLT

>KH.L57.23.v2.A.ND1-1

MTDNGLVYLCAADKEFGRRIPYLFLEELKKQFCSSGSLLQRTTGASAFEFNRDFRNILSGIMDDFNKGKGDQLSTMQNQVGEVTGIMRQNIEKVIERGDKLDDLVDKTEDLQAGAATFKVTAKRIQRKYFWQNKKMLIIIIVIVLIIITLIVLFATGTI

>KH.C8.46.v1.A.ND1-1

MTRYNKGKNTERVALLDDYESDDDFFLKGPSSASDKVKRVQSQVNEVVDVMQTNIGKVLERGDKLEDLQDKSESLADSATHFNTHATRLRKKMWWKDMRTKIIIGVLLLMILIIIIVSIAVKNKGS

>VAMP1-Hs

MSAPAQPPAEGTEGTAPGGGPPGPPPNMTSNRRLQQTQAQVEEVVDIIRVNVDKVLERDQKLSELDDRADALQAGASQFESSAAKLKRKYWWKNCKMMIMLGAICAIIVVVIVIYFFT

>VAMP2-Hs

MSATAATAPPAAPAGEGGPPAPPPNLTSNRRLQQTQAQVDEVVDIMRVNVDKVLERDQKLSELDDRADALQAGASQFETSAAKLKRKYWWKNLKMMIILGVICAIILIIIIVYFST

>VAMP3-Hs

MSTGPTAATGSNRRLQQTQNQVDEVVDIMRVNVDKVLERDQKLSELDDRADALQAGASQFETSAAKLKRKYWWKNCKMWAIGITVLVIFIIIIIVWVVSS

>VAMP4-Hs

MPPKFKRHLNDDDVTGSVKSERRNLLEDDSDEEEDFFLRGPSGPRFGPRNDKIKHVQNQVDEVIDVMQENITKVIERGERLDELQDKSESLSDNATAFSNRSKQLRRQMWWRGCKIKAIMALVAAILLLVIIILIVMKYRT

>VAMP5-Hs

MAGIELERCQQQANEVTEIMRNNFGKVLERGVKLAELQQRSDQLLDMSSTFNKTTQNLAQKKCWENIRYRICVGLVVVGVLLIILIVLLVVFLPQSSDSSSAPRTQDAGIASGPGN

>YKT6-Hs

MKLYSLSVLYKGEAKVVLLKAAYDVSSFSFFQRSSVQEFMTFTSQLIVERSSKGTRASVKEQDYLCHVYVRNDSLAGVVIADNEYPSRVAFTLLEKVLDEFSKQVDRIDWPVGSPATIHYPALDGHLSRYQNPREADPMTKVQAELDETKIILHNTMESLLERGEKLDDLVSKSEVLGTQSKAFYKTARKQNSCCAIM

>VAMP7-Hs

MAILFAVVARGTTILAKHAWCGGNFLEVTEQILAKIPSENNKLTYSHGNYLFHYICQDRIVYLCITDDDFERSRAFNFLNEIKKRFQTTYGSRAQTALPYAMNSEFSSVLAAQLKHHSENKGLDKVMETQAQVDELKGIMVRNIDLVAQRGERLELLIDKTENLVDSSVTFKTTSRNLARAMCMKNLKLTIIIIIVSIVFIYIIVSPLCGGFTWPSCVKK

>VAMP8-Hs

MEEASEGGGNDRVRNLQSEVEGVKNIMTQNVERILARGENLEHLRNKTEDLEATSEHFKTTSQKVARKFWWKNVKMIVLICVIVFIIILFIVLFATGAFS

>SNAP25-Hs

MAEDADMRNELEEMQRRADQLADESLESTRRMLQLVEESKDAGIRTLVMLDEQGEQLERIEEGMDQINKDMKEAEKNLTDLGKFCGLCVCPCNKLKSSDAYKKAWGNNQDGVVASQPARVVDEREQMAISGGFIRRVTNDARENEMDENLEQVSGIIGNLRHMALDMGNEIDTQNRQIDRIMEKADSNKTRIDEANQRATKMLGSG

**Sequences for NCS1 phylogenetic tree:**

>KH.C1.1067.v1.A.SL1-1 (NCS1)

MGKSGSKLSAEDAQTLQTTTHFDKKEIQKWYRDFMKDCPNGYLKKEEFQKVYQQFFPKGNPSKFANFVFNVFDSDKDGFITFKEFISALSVTSRGNLDEKLDWAFNLYDLDHDGFITREEMLNIVDAIYSMVGNAMDLPEDENTPEKRVNKIFCQMDQNKDGKLTKDEFREGSKCDPYIVKALSAGLGGAESCPS

>KH.C9.113.v1.A.ND1-1
MGNQESTQSLSEHEIARLSAETNFTPAEVAQWYKGFKRDCPSGKFTMARFQAIAEGFFPAGDAENFTKFIFNGMDLEEDASLNFESFIKVTSLLARGTKEEKLKWAFHVRDIDHTGFITKEKMQLVEDSVFSMIANHVPLPDDENTAEKRTEKLYNLIEKDQDGKVTVDQFKVGLNQDRDIVNALSLHDTVQ

>KH.S534.4.v2.A.SL3-1
MGCAKSKTRDEDCKQLESLTNFTPEELKKCYDDFQQESTSGKMDRTKFDEFYKKFFNRDPRFVDHLFRTFDFNNDGFINFREFVCGLSITTRGTPEEKLTWTFNVYDVNNDGTITRDEMLQIMRAIYAMNGISEPEQLKSGSDAFEGLDSNGDGLISVAEFVKGVKRDERLLEFLQRTIDVQQK

>KH.S534.6.v1.A.nonSL2-1
MGQQHPSLKPRMLDDLMRMSDFSADELKIWYNEFSKDSVSGFLSKNDFIKIYKGLFPHGDASGFADHVFRTFDANKDGVLNFREFVIGLSLTMKGPLDDKLYWAFKLYDVDGNGFVTKDEMNEIIAVSNAIFTVCCTDTTDDDVIMSGREAFKTMDTDEDGKVSWGEFRKGIHSNRVFMLIVEAKMKNAGVTSR

>KH.C8.342.v1.A.SL2-1
MGKQNSKLKPEDIRDLRAATEFNEHELQEWYKGFIKDCPSGNLTMDEFQKIYANFFPQGDASKFAAHVFRTFDSNGDSTIDFREFIVALSVTSRGNLEEKLKWAFSMYDCDGNGIISRDEMLEIVRAIYKMVGAVMKMPEDESTPEKRTDKIFKQMDKNLDGSISLEEFVEGAKKDPSIVRLLQCTPGSGGLGM

>KH.C9.274.v1.A.SL1-1

MGNRKSKLKPEVLEKLTKQTKFTEAELHQWHKGFLHDCPTGKLSYEEFQGIYRQFFPQGNSAKFAKLVFTTFDDNKDGTVEFEEFIIALSVTSRGTLDEKLHWAFQLYDLDNDGFITKDEMLNIVEAIFAMVGDAVNLPAEENTPQKRVEKIFKVMDKNKDGKLTKDEFLVGAKSDPSIVQALSIYDGLV

>NCS1-Hs

MGKSNSKLKPEVVEELTRKTYFTEKEVQQWYKGFIKDCPSGQLDAAGFQKIYKQFFPFGDPTKFATFVFNVFDENKDGRIEFSEFIQALSVTSRGTLDEKLRWAFKLYDLDNDGYITRNEMLDIVDAIYQMVGNTVELPEEENTPEKRVDRIFAMMDKNADGKLTLQEFQEGSKADPSIVQALSLYDGLV

>NCALD-Hs

MGKQNSKLRPEVMQDLLESTDFTEHEIQEWYKGFLRDCPSGHLSMEEFKKIYGNFFPYGDASKFAEHVFRTFDANGDGTIDFREFIIALSVTSRGKLEQKLKWAFSMYDLDGNGYISKAEMLEIVQAIYKMVSSVMKMPEDESTPEKRTEKIFRQMDTNRDGKLSLEEFIRGAKSDPSIVRLLQCDPSSAGQF

>HPCA-Hs

MGKQNSKLRPEMLQDLRENTEFSELELQEWYKGFLKDCPTGILNVDEFKKIYANFFPYGDASKFAEHVFRTFDTNSDGTIDFREFIIALSVTSRGRLEQKLMWAFSMYDLDGNGYISREEMLEIVQAIYKMVSSVMKMPEDESTPEKRTEKIFRQMDTNNDGKLSLEEFIRGAKSDPSIVRLLQCDPSSASQF

>HPCAL4-Hs

MGKTNSKLAPEVLEDLVQNTEFSEQELKQWYKGFLKDCPSGILNLEEFQQLYIKFFPYGDASKFAQHAFRTFDKNGDGTIDFREFICALSVTSRGSFEQKLNWAFEMYDLDGDGRITRLEMLEIIEAIYKMVGTVIMMRMNQDGLTPQQRVDKIFKKMDQDKDDQITLEEFKEAAKSDPSIVLLLQCDMQK

>VSNL1-Hs

MGKQNSKLAPEVMEDLVKSTEFNEHELKQWYKGFLKDCPSGRLNLEEFQQLYVKFFPYGDASKFAQHAFRTFDKNGDGTIDFREFICALSITSRGSFEQKLNWAFNMYDLDGDGKITRVEMLEIIEAIYKMVGTVIMMKMNEDGLTPEQRVDKIFSKMDKNKDDQITLDEFKEAAKSDPSIVLLLQCDIQK

>Frq1-Dm

MGKKSSKLKQDTIDRLTTDTYFTEKEIRQWHKGFLKDCPNGLLTEQGFIKIYKQFFPQGDPSKFASLVFRVFDENNDGSIEFEEFIRALSVTSKGNLDEKLQWAFRLYDVDNDGYITREEMYNIVDAIYQMVGQQPQSEDENTPQKRVDKIFDQMDKNHDGKLTLEEFREGSKADPRIVQALSLGGG

>Frq2-Dm

MGKKNSKLKQDTIDRLTTDTYFTEKEIRQWHKGFLKDCPNGLLTEQGFIKIYKQFFPDGDPSKFASLVFRVFDENNDGAIEFEEFIRALSITSRGNLDEKLHWAFRLYDVDNDGYITREEMYNIVDAIYQMVGQQPQTEDENTPQKRVDKIFDQMDKNHDDRLTLEEFREGSKADPRIVQALSLGGD

>Efcab11-Cr (KH.C10.577.v1.A.ND1-1)

MAVSLRRLQTIFNHCDEDKKGFLNREDLKVAMLILFGYKPSKYEIQQLLDDGEGNKKLMNFETFKSIMAASKHDPDHEIRDIFRMFDTHCRGFLIFDDVKKAFHAVAPHISDATIRACCEEMCRESDGRISYREFAEAMRHGHVHMEHHENYDHFFNSNVCFS

**Sequences for NARS1 phylogenetic tree:**

>KH.C12.45.v1.A.SL1-1 (NARS1)

MGDSVEQQVNKLSIGELYTSEKTGCDESGDGTEAKPFKTILRAMMFHQSEPFPTLYVDSKEPNKKFDPIAKAQLKKQTKIFTAEKRKDVARKVREQEDAERREKNLEDAKKIIISEDKSLPAAKSIKIGAATAHRGQRVKVNGWSHRIRRQGKTLMFIVLRDGSGYLQAVLNDQLCQTYNAVMLSTESTVCLYGVITPVMEGKSAPGGHELICDYWELIGLAPPGGIDTVLNQEADVDVMLDNRHLVIRGEQTSKILRVRSYLMQAFRDHYFERGYYEVTPPCLVQTQCEGGSTLFKLDFFGEQAYLTQSSQLYLETCLPSLGDVFCLAESYRAEQSRTRRHLAEYTHVEAECAFITFNDLLDRLEDLICDVVDKLLKSPAGELIKDLNPGFVAPKRPFRRMDYAEAIVYLKEHDIKKEDGSYYEFGEDIPEMPERKMTDQINEPILLCRFPYKIKSFYMQKCKDNLELTESVDLLMPNVGEIVGGSMRMDNHEELLEGYKSEGVDPSKYYWYTDQRKFGTCEHGGYGLGLERLLTWLTDRYHIRDVCLYPRFLERCCP

>KH.C7.539.v1.A.SL1-1

MADVENVPLGDDGKPMSKKALKKQQKDAEKAAKKAQRQQEQADAQQKADADDVSKDLYGDMVMIQSSEKPDRCLQPIGDIGPNLAEKNIWVRGRLHTSRGTGKQCFTVLRHQQSTIQALIFVGEKISKQMVKFCANISKESIVDVYGVVKKVEQKVESCTQDDVEIHIGKIFVVSKAAPQLPLLIEDASRPELEDGKNEEGGRPRVNLDTRLDNRILDLRTTTKQAIFRLEAGVCKLFRDTLTAKGFCEIHTPKIISAASEGGANVFTVSYFKTNAFLAQSPQLYKQMAIAADFEKVFTVGAVFRAEDSNTHRHLTEFVGLDLEMAFQYHYHEVLQVIGDMFISIFKGLRESYQKEITTVARQYPAEPFKFLEPTLILQYPEAVKMLNDAGVEMQDNEDLSTPSEKLLGRLVRQKYDTDYYILDKYPLEVRPFYTMPDPDNPKWSNSYDMFMRGEEILSGAQRIHDPEFLSERAKAHGIDLKTIEAYIDSFKYGCPPHAGGGIGMERVVMLYLGLHNIRSSSLFPRDPKRLTP

>KH.C2.670.v1.A.ND1-1

MFRLCKLSYSTVTTALKRFKIKDVVNGEAVSQNGWIQGWVQSIRKHKNFMFVDIVDGTSLNPLQVVLPSHMFNSQLQNGAAVSANGTFVETDLNVNKIEMVCEEIQVVGPCDPETYPFLKHTPHDMQHLRSYPHLRMRRTSMINAFKLRNQLEMLFHEFFQNEGFTHVHTPIITLSDCEGAGELFSIKSDTSTNTDMENSENHFFNKPAYLTVSGQMHLEACAMSLGSVYTLSPVFRAEHGISRKHLSEFRMLEVEVAFTQDLHEILDLIEDSIKFCIERVQSVPLKKFDDGYKDIPPQHHKAVKNATCSNYVRIPYSEALEILQKNSHKLTTAAPVWGDDFNTDQESFLVKHFGLTPVFIINFPKVAKPFYMYANNDNNTVAAVDLIFPFCGEICGGSLREHRLELLQDSLRTSGLNEKDYEWYLDLRRFGSAPHGGYGLGFDRLVHFLLGTRNIKDAVPFPRTPHSCPL

>KH.C3.437.v2.A.SL1-1

MADNGDCEEKKISKNEMKRQLKAQQKAKEKAEKAAKALENAPKEVKPEKTASKEEETLDPNQYLQIRKNTITTLRQNNIEPYPHKFHVSISLSDYVEKYNNIEVGSHLNDQQVSIAGRIHAKREAGPKLIFYDVRGDGVKLQVMANSKMYSSEEAYQEINERTRRGDIIGVIGHPAKTKKGELSIVPNTIEILSPCLHMLPHLHFGLKDKETRYRQRYLDLIMNDQTRQKFITRAKIISYIRSFFDQMGFLEVETPMMNMVAGGATAKPFITHHNDLDMDLYMRVAPELYLKMLVVGGLDRVYEIGRLFRNEGIDMTHNPEFTSCEFYMAYADYEDLMKISETLISGMVKQICGSYKLTYHPDGPDSEGYEVDYTPPFRRLRMLPDLERLTGMEFPKPTELHTAGAQARLDEICVKLGVECPPPRTTARLLDKLVGDYLEVNCINPTFITEHPEIMSPLAKWHRSIKGLTERFELFVNKKEICNAYTELNDPMIQRQRFEQQALDKAAGDDEAQMVDENFCTALEYGLPPTGGWGMGIDRLTMFLTDSNNIKEVLLFPAM

KPDDQVPKKAEDSPST

>KH.C11.248.v1.A.SL2-1

MLKLFRVVRPILFKFRPKFNPVTICIVTKQFTSNSAIPQVVQSIKNEKLPNSKFSYRSHTCGELRIKNAGEAVSLCGWLEYKRKDFLVLRDSFGSVQILITGYEDKNQTFQNVKPESVVKVTGKVTTRPEGQANPNMITGEIEIIPDNIEILNKCKFLPFRVQKFAGKGERIRQKYRYLDIRGDILQQNLRLRSKVLSKMRKSLEENCFVEVETPTLFHPTPGGAREFIVPSSSNPGKFYSLPQSPQQLKQLLMIGGVDRYYQFARCYRDEPFKRDRQVEFTQLDIEMSFVDVNDVIKLSEEVIKSSWPRKFSDESFKLITYDDAMSKYGSDKPDTRFGMLLHDITTFIPDTLVELFQGDHSDPVNIQAVVVENAQKHFSSKFRRYIGSRMTSAIDDPSPYALLGLVNGQWSGYHKNTNIGSMVTKQLVDHLQLGGSDIVVVSWGEKRSAQKTLGALRRIISRIFDEISVKHRNVDLDQFLWVVDFPLFEADPYTGSLCTVHHPFTSPHPDDVHLLEVDPLKVRSQAYDLVLNGNEVGGGSIRIHDSKLQRQIFTLLNLD

FTELSYLLEALNSGAPPHGGIAFGIDRIMSILCGTETIRDVIAFPKNSRGEDLMTNSPISINSNILKEFHIKTNVSDK

>YARS1-Hs

MGDAPSPEEKLHLITRNLQEVLGEEKLKEILKERELKIYWGTATTGKPHVAYFVPMSKIADFLKAGCEVTILFADLHAYLDNMKAPWELLELRVSYYENVIKAMLESIGVPLEKLKFIKGTDYQLSKEYTLDVYRLSSVVTQHDSKKAGAEVVKQVEHPLLSGLLYPGLQALDEEYLKVDAQFGGIDQRKIFTFAEKYLPALGYSKRVHLMNPMVPGLTGSKMSSSEEESKIDLLDRKEDVKKKLKKAFCEPGNVENNGVLSFIKHVLFPLKSEFVILRDEKWGGNKTYTAYVDLEKDFAAEVVHPGDLKNSVEVALNKLLDPIREKFNTPALKKLASAAYPDPSKQKPMAKGPAKNSEPEEVIPSRLDIRVGKIITVEKHPDADSLYVEKIDVGEAEPRTVVSGLVQFVPKEELQDRLV

VVLCNLKPQKMRGVESQGMLLCASIEGINRQVEPLDPPAGSAPGEHVFVKGYEKGQPDEELKPKKKVFEKLQADFKISEECIAQWKQTNFMTKLGSISCKSLKGGNIS

>DARS1-Hs

MPSASASRKSQEKPREIMDAAEDYAKERYGISSMIQSQEKPDRVLVRVRDLTIQKADEVVWVRARVHTSRAKGKQCFLVLRQQQFNVQALVAVGDHASKQMVKFAANINKESIVDVEGVVRKVNQKIGSCTQQDVELHVQKIYVISLAEPRLPLQLDDAVRPEAEGEEEGRATVNQDTRLDNRVIDLRTSTSQAVFRLQSGICHLFRETLINKGFVEIQTPKIISAASEGGANVFTVSYFKNNAYLAQSPQLYKQMCICADFEKVFSIGPVFRAEDSNTHRHLTEFVGLDIEMAFNYHYHEVMEEIADTMVQIFKGLQERFQTEIQTVNKQFPCEPFKFLEPTLRLEYCEALAMLREAGVEMGDEDDLSTPNEKLLGHLVKEKYDTDFYILDKYPLAVRPFYTMPDPRNPKQSNSYDMFM

RGEEILSGAQRIHDPQLLTERALHHGIDLEKIKAYIDSFRFGAPPHAGGGIGLERVTMLFLGLHNVRQTSMFPRDPKRLTP

>DARS2-Hs

MYFPSWLSQLYRGLSRPIRRTTQPIWGSLYRSLLQSSQRRIPEFSSFVVRTNTCGELRSSHLGQEVTLCGWIQYRRQNTFLVLRDFDGLVQVIIPQDESAASVKKILCEAPVESVVQVSGTVISRPAGQENPKMPTGEIEIKVKTAELLNACKKLPFEIKNFVKKTEALRLQYRYLDLRSFQMQYNLRLRSQMVMKMREYLCNLHGFVDIETPTLFKRTPGGAKEFLVPSREPGKFYSLPQSPQQFKQLLMVGGLDRYFQVARCYRDEGSRPDRQPEFTQIDIEMSFVDQTGIQSLIEGLLQYSWPNDKDPVVVPFPTMTFAEVLATYGTDKPDTRFGMKIIDISDVFRNTEIGFLQDALSKPHGTVKAICIPEGAKYLKRKDIESIRNFAADHFNQEILPVFLNANRNWNSPVANFIME

SQRLELIRLMETQEEDVVLLTAGEHNKACSLLGKLRLECADLLETRGVVLRDPTLFSFLWVVDFPLFLPKEENPRELESAHHPFTAPHPSDIHLLYTEPKKARSQHYDLVLNGNEIGGGSIRIHNAELQRYILATLLKEDVKMLSHLLQALDYGAPPHGGIALGLDRLICLVTGSPSIRDVIAFPKSFRGHDLMSNTPDSVPPEELKPYHIRVSKPTDSKAERAH

>NARS1-Hs

MVLAELYVSDREGSDATGDGTKEKPFKTGLKALMTVGKEPFPTIYVDSQKENERWNVISKSQLKNIKKMWHREQMKSESREKKEAEDSLRREKNLEEAKKITIKNDPSLPEPKCVKIGALEGYRGQRVKVFGWVHRLRRQGKNLMFLVLRDGTGYLQCVLADELCQCYNGVLLSTESSVAVYGMLNLTPKGKQAPGGHELSCDFWELIGLAPAGGADNLINEESDVDVQLNNRHMMIRGENMSKILKARSMVTRCFRDHFFDRGYYEVTPPTLVQTQVEGGATLFKLDYFGEEAFLTQSSQLYLETCLPALGDVFCIAQSYRAEQSRTRRHLAEYTHVEAECPFLTFDDLLNRLEDLVCDVVDRILKSPAGSIVHELNPNFQPPKRPFKRMNYSDAIVWLKEHDVKKEDGTFYEFGEDIP

EAPERLMTDTINEPILLCRFPVEIKSFYMQRCPEDSRLTESVDVLMPNVGEIVGGSMRIFDSEEILAGYKREGIDPTPYYWYTDQRKYGTCPHGGYGLGLERFLTWILNRYHIRDVCLYP

RFVQRCTP

>NARS2-Hs

MLGVRCLLRSVRFCSSAPFPKHKPSAKLSVRDALGAQNASGERIKIQGWIRSVRSQKEVLFLHVNDGSSLESLQVVADSGLDSRELNFGSSVEVQGQLIKSPSKRQNVELKAEKIKVIGNCDAKDFPIKYKERHPLEYLRQYPHFRCRTNVLGSILRIRSEATAAIHSFFKDSGFVHIHTPIITSNDSEGAGELFQLEPSGKLKVPEENFFNVPAFLTVSGQLHLEVMSGAFTQVFTFGPTFRAENSQSRRHLAEFYMIEAEISFVDSLQDLMQVIEELFKATTMMVLSKCPEDVELCHKFIAPGQKDRLEHMLKNNFLIISYTEAVEILKQASQNFTFTPEWGADLRTEHEKYLVKHCGNIPVFVINYPLTLKPFYMRDNEDGPQHTVAAVDLLVPGVGELFGGGLREERYHFLEERLA

RSGLTEVYQWYLDLRRFGSVPHGGFGMGFERYLQCILGVDNIKDVIPFPRFPHSCLL

>KARS1-Hs

MAAVQAAEVKVDGSEPKLSKNELKRRLKAEKKVAEKEAKQKELSEKQLSQATAAATNHTTDNGVGPEEESVDPNQYYKIRSQAIHQLKVNGEDPYPHKFHVDISLTDFIQKYSHLQPGDHLTDITLKVAGRIHAKRASGGKLIFYDLRGEGVKLQVMANSRNYKSEEEFIHINNKLRRGDIIGVQGNPGKTKKGELSIIPYEITLLSPCLHMLPHLHFGLKDKETRYRQRYLDLILNDFVRQKFIIRSKIITYIRSFLDELGFLEIETPMMNIIPGGAVAKPFITYHNELDMNLYMRIAPELYHKMLVVGGIDRVYEIGRQFRNEGIDLTHNPEFTTCEFYMAYADYHDLMEITEKMVSGMVKHITGSYKVTYHPDGPEGQAYDVDFTPPFRRINMVEELEKALGMKLPETNLFETEETR

KILDDICVAKAVECPPPRTTARLLDKLVGEFLEVTCINPTFICDHPQIMSPLAKWHRSKEGLTERFELFVMKKEICNAYTELNDPMRQRQLFEEQAKAKAAGDDEAMFIDENFCTALEYGLPPTAGWGMGIDRVAMFLTDSNNIKEVLLFPAMKPEDKKENVATTDTLESTTVGTSV

Cr: Ciona robusta

Hs: Homo sapiens

Dm: Drosophila melanogaster
